# MTQuant: “Seeing” Beyond the Diffraction Limit in Fluorescence Images to Quantify Neuronal Microtubule Organization

**DOI:** 10.1101/074047

**Authors:** Roshni Cooper, Shaul Yogev, Kang Shen, Mark Horowitz

**Affiliations:** Department of Electrical Engineering, Stanford University, Stanford, CA 94305, USA; Department of Biology, Howard Hughes Medical Institute, Stanford University, Stanford, CA 94305, USA.

## Abstract

**Motivation:** Microtubules (MTs) are polarized polymers that are critical for cell structure and axonal transport. They form a bundle in neurons, but beyond that, their organization is relatively unstudied.

**Results:** We present MTQuant, a method for quantifying MT organization using light microscopy, which distills three parameters from MT images: the spacing of MT minus-ends, their average length, and the average number of MTs in a cross-section of the bundle. This method allows for robust and rapid *in vivo* analysis of MTs, rendering it more practical and more widely applicable than commonly-used electron microscopy reconstructions. MTQuant was successfully validated with three ground truth data sets and applied to over 3000 images of MTs in a *C. elegans* motor neuron.

**Availability:** MATLAB code is available at http://roscoope.github.io/MTQuant

**Supplementary information:** Supplementary data are available at *Bioinformatics* online.

## 1 Introduction

Microtubules (MTs) are dynamic polymers that are formed by polymerization of alpha-beta tubulin dimers. MTs play critical roles in cell architectures and as tracks for intracellular transport. The complex cellular morphology of neurons, and the long distance intracellular transport that occurs in them, rely heavily on proper MT function (Desai and Mitchison, 1997; Kapitein and Hoogenraad, 2015; Conde and Caceres, 2009). This is evident by the large number of neurodegenerative diseases that are linked to mutations that disrupt MT structure or MT dependent cargo transport (Tischfield *et al*., 2010; Millecamps and Julien, 2013).

In axons, MTs are tiled side by side and head to tail to form a bundle (Kapitein and Hoogenraad, 2015). Neuronal MTs have mostly been characterized with respect to their dynamic behavior (Applegate *et al*., 2011; Demchouk *et al*., 2011) and polarity, as these properties can be monitored relatively easily by fluorescence-tagging of a plus-end binding protein (Stepanova *et al*., 2003). However, snapshots of MT dynamics do not reveal the steady-state size of the polymer, which is the outcome of growth and shrinkage, nor the number of polymers or their spacing, i.e., the axial distance between the starting locations of consecutive MTs (see Supp. Table 1). To obtain these parameters, which we collectively refer to as MT organization, it is necessary to perform serial reconstruction of electron microscopy sections, a task which is extremely slow, laborious, and error-prone. The MT diameter (~24nm) and the tight bundling of neuronal MTs make the analysis of neuronal MT organization challenging even with super-resolution methods, although promising results have recently been reported (Mikhaylova *et al*., 2015; Balint *et al*., 2013).

We developed a rapid and robust fluorescence-based method for the assessment of MT organization in neuronal processes, titled MTQuant, and applied it in the *C. elegans* motor neuron DA9 (Yogev *et al*., 2016b). The method uses a GFP::alpha-tubulin transgene to label MTs and then applies image processing and optimization techniques to the total bundle intensity, thereby enabling the extraction of parameters of MT organization. Here, we describe in detail the image processing algorithms that are used in the analysis of the fluorescent signal. We also propose improvements to our original algorithms, which are meant to increase the accuracy of the method. Finally, we apply a procedure that refines the distribution of the organization parameters. MTQuant has been validated against three data sets, including mathematical simulations and two types of fluorescent MT images. This framework allows for higher-level comparison of MT organization between genotypes, developmental time-points, and drug treatments, and can therefore greatly enhance our understanding of neuronal MTs.

## 2 Approach

The MTQuant framework is outlined in Algorithm 1 and depicted in the block diagram in Supp. Fig. 1. Each step of MTQuant is detailed in the Methods section. MTQuant allows biologists to begin with a confocal image of the entire MT bundle, and ultimately extract the spacing, coverage, and length of the MTs of a population of animals. The framework consists of two main steps: the analysis of individual worms, followed by refinement of the distributions of the organization parameters. The individual worm analysis relies on the fact that MTs form a bundle neurons, as shown in Fig. 1(a). It uses images of MTs labeled with two fluorescent proteins: GFP labels the alpha tubulin protein that is part of the alpha-beta dimer which polymerizes to form MTs, while TagRFP specifically binds to patronin, a protein present at one tip of each microtubule, called the minus-end. The punctate red channel in Fig. 1(b) shows the red patronin staining of the minus-ends of the MTs.

**Fig. 1.**
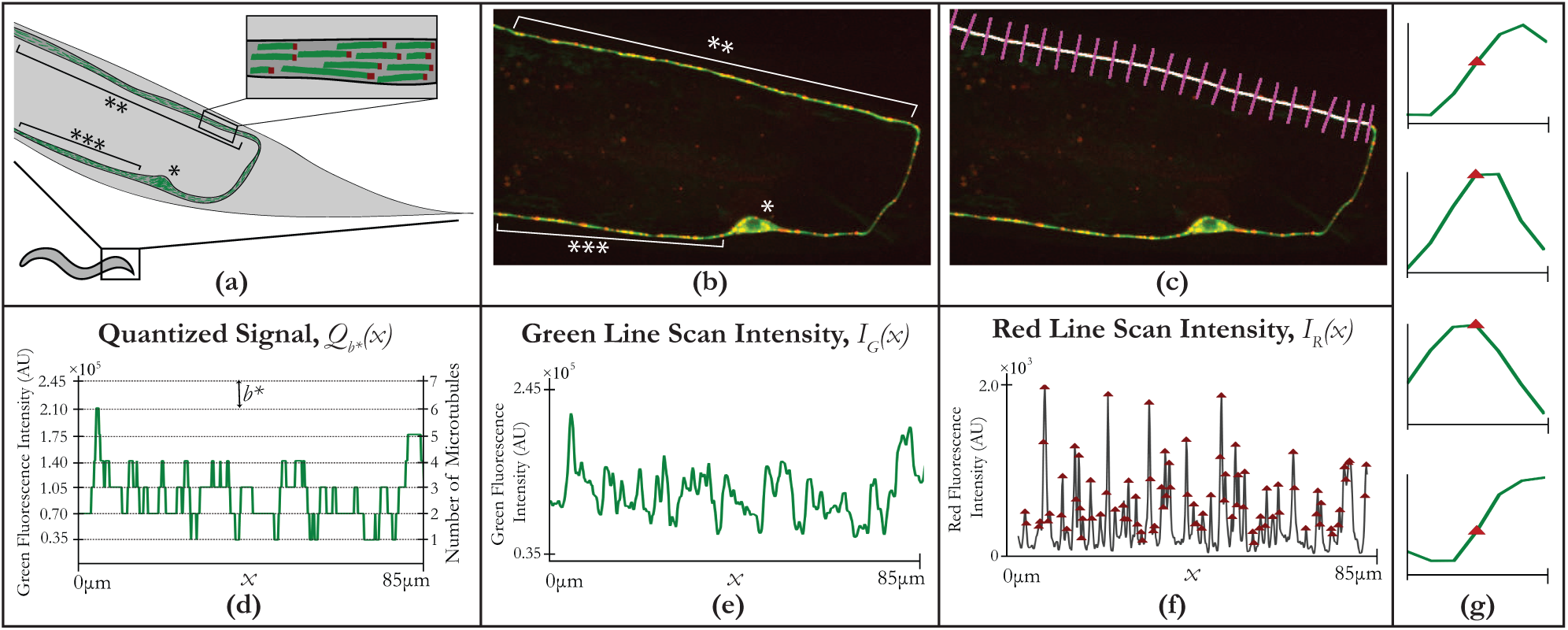
Neuron Tracing. (a) Illustration of DA9 in the tail of C. elegans. The inset shows the bundling of microtubules side-by-side and end-to-end. In (a) and (b), the (*) is the cell body, (**) is the axon, and (***) the dendrite. (b) Fluorescence image of MT bundle in DA9 neuron. Green channel shows microtubules and red channel shows MT minus-ends. (c) Line scan extraction. White line forms spline which traces neurite of interest. Magenta lines are subset of line segments perpendicular to spline. Image intensity is summed along magenta segments beginning closest to cell body (right) to form line scan in (e). (d) Expected MT bundle intensity, *Q*_*b*_*(*x*). (e) Line scan, or trace of green tubulin intensity, *I*_*G*_(*x*). *I*_*G*_(*x*) can be quantized by the intensity contribution from one MT, called the single MT brightness, *b*\*, to estimate the underlying MT bundle intensity, *Q*_*b*_* (*x*) in (d). (f) Red patronin signal trace along neurite. Red triangles represent detected MT minus-ends. (g) Average green tubulin intensity around MT minus-ends for four animals. Segments of green signal around all MT minus-ends in a single animal were aligned and averaged. MT minus-ends (red triangles), on average, align with an increase in the green signal.

The framework traces the green tubulin intensity and the red patronin intensity along the white line in Fig. 1(c) to yield the green and red intensity line scans shown in Fig. 1(e)-(f), respectively. Axonal MTs are uniformly oriented with their minus-ends pointing toward the cell body. Hence, the red dots in Fig. 1(a)-(b), are all on the same side of the MTs, and, when traversing the axon away from the cell body, the location of a red puncta, on average, corresponds to a increase in the green intensity trace, as shown in Fig. 1(g).

We expect the green tubulin intensity to be a quantized function, as in Fig. 1(d), where each MT contributes to an additional level of intensity. We assume that the intensity of each MT in an animal is the same, and, furthermore, constant along the neurite. Hence, a change in the level of the quantized signal, i.e., a step up or a step down, marks the beginning or the end of a MT, respectively. In reality, however, the imaging system blurs the signal with its point spread function and adds noise to the signal, resulting in an intensity profile such as the one in Fig. 1(e). We assume the noise in the system is zero-mean, so quantizing the green tubulin line scan with the correct intensity contribution from a single MT, *b*\* in Fig. 1(d), yields a reasonable approximation of the underlying MTs. We use the following notation when describing the rounding: ⌊*x*⌋ rounds *x* to the nearest integer.

Correctly identifying the intensity of a single MT, referred to as the single MT brightness, is critical to the success of MTQuant, and is described in detail in Section 3.2. The single MT brightness must be recalculated for each animal, as the imaging position and the fluorescent protein expression varies from animal to animal. Another important piece of information MTQuant uses is the number of microtubules in the neuron. That number is, of course, closely related to the number of peaks in the intensity of the red line scan. Since relatively little is known about the MT minus-end locations, we assume the MT minus-ends are distributed independently, according to a Poisson process (Bertsekas and Tsitsiklis, 2002). Section 3.1.3 will detail how we use the red scan to estimate the locations of the MT minus-ends.

Armed with the single MT brightness and the MT minus-ends, we can calculate organization parameters, as shown in Supp. Table 1. The spacing of the MTs is simply the average distance between the MT minus-ends. If we quantize the traced intensity signal with the single MT brightness, the value of the quantized function is the number of microtubules in the axon cross-section at any point along the neuron. Then average coverage is the average value of the quantized signal. The average MT length is slightly less straightforward since we cannot identify individual MTs. It is the integral of the quantized signal divided by the total number of MTs. Section 3.3 will describe all of these parameter calculations in more detail.

### Algorithm 1: MTQuant Overview

1. Analyze Individual Worms For each worm:
  a. Trace neurite to extract green & red line intensities
  b. Identify “single MT brightness” using grid-search optimization
  c. Calculate spacing, coverage, and length of MTs
2. Refine distributions of organization parameters with expectation-maximization

Finally, after calculating the organization parameters for each individual animal, MTQuant aggregates the values of each parameter for a group of worms, i.e. all worms of the same age, genotype, etc., to remove some of the measurement noise that affects each calculation. In this step, expectation-maximization is used to refine the distributions of the organization parameters.

An initial version of MTQuant was applied to the data in (Yogev *et al*., 2016b,a). The framework described here, and outlined in Algorithm 1, builds on the initial algorithm used in those papers by improving the accuracy of the calculation of the singleMTbrightness and by implementing an additional distribution refinement following the analysis of the individual animals to generate more accurate distributions of spacing, coverage, and length across a population of animals.

## 3 Methods

The extraction of the microtubule organization parameters from fluorescence images is illustrated in the block diagram in Supp. Fig. 1. Each animal is analyzed independently, and then the organization parameters of a group of worms are aggregated to attain a more refined distribution of the parameters. The individual worm analysis involves image processing to extract the intensity of the microtubule bundle, followed by the identification of the brightness of a single microtubule, and finally the calculation of the organization parameters. Each block in this pipeline, as well as a description of a microtubule simulation method used to verify the accuracy of MTQuant, is described below.

### 3.1 Image Processing

We begin the analysis of each individual animal by segmenting the neurite of interest and then tracing the intensity along it. This initial process involves aligning the two color channels and asking the biologist to select which section of the neuron to analyze, i.e., the dendrite or the axon. Then we identify the spline that traces the neuron most closely and interpolate the pixel values along the spline to extract the intensity of the entire microtubule bundle from the green image. Finally, we use this same extracted spline to trace the intensity in the red image and identify the peaks in the red signal, which correspond to the starting locations of the microtubules. In the following sections, we use the green intensity and the red peak locations to continue the process of extracting the organization parameters of the microtubules.

#### 3.1.1 Preprocessing

MTQuant begins by aligning the green color channel with the red color channel for each worm. The alignment uses correlation to find the best matching translation of the green and red channels to account for misalignment due to the imaging system. The algorithm searches for the best correlation in a small window around the origin. In the case that a small, simple-to-find translation is an insufficient transformation, the animal was moving during imaging and is too misaligned to use for this algorithm. This severe misalignment happens infrequently because the animals are anesthetized.

A graphical user interface (GUI) asks the user to select a few points along the neurite of interest. This GUI allows the user to select, for example, only the dendrite, or a specific section of the axon, as opposed to automatically tracing the entire neuron. The points clicked by the user are connected linearly and the lines are dilated to form the “neurite mask.” The user also identifies the orientation of the worm, i.e., where the minus-ends are located, and selects a few points inside the worm from which to identify the background autofluorescence intensity of the animal, which is subtracted from the observed intensities.

#### 3.1.2 Line Scan Extraction

Then the algorithm iteratively adjusts the mask to identify the exact pixels containing the neuron. See Algorithm 2. The algorithm iteratively fits a spline to the current neuron estimate, and searches the line segments perpendicular to the spline to find the brightest pixel that is also close to the spline. Then we update the spline to include this brightest point. We penalize any new points that are too far from the spline, in order to avoid readjusting the spline to include bright background noise or autofluorescence from other tissues. This regularization is controlled using the parameter *α* in Step 2(a)ii of Algorithm 2. We selected *α* based on visual experiments. At the end of each iteration, we re-space the points by interpolating the points along the new spline and selecting points that are separated by exactly zero or one pixel in each direction. This re-spacing ensures that every pixel in the neuron is traced but also avoids fitting an overly-precise spline that provides no more accuracy but increases computation time. The algorithm iterates until the spline is no longer changing significantly. An example image and spline are shown in Fig. 1(b)-(c).

The next step is to trace the signal intensity along the spline. Due to the blurring of the point spread function of the microscope and the positioning of the MT bundle, the intensity of the fluorescence markers extends over several pixels perpendicular to the neuron. For every pixel along the spline of the neurite, the intensities of the interpolated points on the line segment perpendicular to the spline are summed, i.e. the magenta segments in Fig. 1(c). The grid is interpolated using linear interpolation. This summation results in a signal referred to as a “line scan.” For the green channel, the line scan represents the total tubulin intensity at any point along the neuron, and for the red channel, the line scan represents the total minus-end intensity at any point along the neuron.

##### Algorithm 2: Identify Brightest & Shortest Path

1. Fit spline *S*^0^ = 〈*x*^0^, *y*^0^〉 to points in user-selected neurite mask
2. Repeat until convergence (i.e., ∥〈*x*^*t*+1^,*y*^*t*+1^〉 − 〈*x*^*t*^,*y*^*t*^〉∥ < ϵ);
  a. For each point 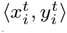 in *S*,
    1. Identify the line segment *P*_*i*_, perpendicular to *S* at 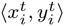
    2. Update 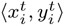 to brightest and nearest point:

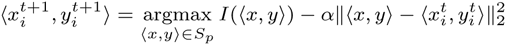
  b. Refit spline to new points 〈*x*^*t*+1^,*y*^*t*+1^〉
  c. Respace the points 〈*x*^*t*+1^,*y*^*t*+1^〉 to be zero or one pixel apart in each direction

To verify that these line scans accurately trace the total intensity along the neuron, we simulated curves of uniform intensity and traced the intensity along them. To simulate these curves, we first simulated a single-pixel curve in a large image (5120×5120 pixels). This curve and resolution simulated the “real” MT bundle. Then the curve was blurred with a Gaussian and binned to simulate the blurring and binning of the pixels in the camera sensor of the imaging system. The resulting image, cropped and shown in Supp. Fig. 2(a), was 512×512 pixels and blurred. When we traced the intensity along the simulated curve, i.e., the red spline in Supp. Fig. 2(b), we saw a uniform intensity along the entire spline using both linear interpolation and cubic interpolation. Supp. Fig. 2(c)-(d) shows the intensity calculated using both linear and cubic interpolation. Cubic interpolation is more computationally intensive than linear interpolation, but, as expected, it resulted in a slightly smoother signal than its linear counterpart (Thévenaz *et al.,* 2009). However, even at varying curvature angles in the image, the maximum error in the linearly interpolated signal is only 1.3% of the average value of the signal. That amount is well below the level of microtubule intensities we would expect to detect, as there are typically between one and twenty microtubules the neuron (Chalfie and Thomson, 1979). Additionally, since the curvature of the neuron is not expected to vary greatly, errors from changes in the slope of the line should be small, even using linear interpolation.

#### 3.1.3 Red Peak Extraction

Before extracting the brightness of a single MT, the locations of the MT minus-ends must be identified. They are extracted from the red line scan in Fig. 1(f). First, the line scan is smoothed with a three-pixel-wide sliding window in order to smooth any extraneous noisy peaks. Then peaks in the red line scan are identified as local maxima, and readjusted as necessary: peaks that are too small compared to neighboring peaks are ignored, and peaks that are particularly wide or tall are considered to be either one MT minus-end or two MT minus-ends using the stochastic method described below. This correction compensates for red minus-ends blurred together by the imaging system’s point spread function. To determine whether to consider a red punctum as one or two minus-ends, a Bernoulli random variable is used. The mean of the random variable is the sum of two probabilities: a height probability and a width probability. The height probability is simply the red intensity value of the peak in question divided by the maximum intensity of the red signal. Hence, taller peaks are more likely to be considered two minus-ends. The width probability is the sigmoid function applied to a measure of the width of the peak: the peak is scaled to be a probability distribution of intensities, and the width measure is calculated as the standard deviation of this probability distribution. Wider peaks have a larger width measure, and using the sigmoid of the width measure rather than the measure itself further encourages wide peaks to be considered two peaks. The mean of the Bernoulli random variable generated by the sum of the height probability and the width probability is typically greater than 0.9, so the random variable is generally true, and most peaks are considered multiples. This dual labeling of the red peaks is consistent with the expectation that minus-ends probably fall close to one another when drawn from a Poisson process.

### 3.2 Single MT Brightness Identification

The calculation of the coverage and the length are predicated on knowing the intensity contribution of a single microtubule, *b*\*, referred to as the single MT brightness. The single MT brightness, *b*\*, needs to be calculated for each worm individually, because *b*\* varies due to various imaging factors, e.g. the distance between the worm and the microscope objective, the amount of tissue between the neuron and the objective, and the extend of the incorporation of fluorophores into the MTs for that specific worm.

Since we observe only the integrated signal of the entire MT bundle, calculating the single MT brightness directly would require solving an impossibly complex optimization with an infinite number of possible solutions. Instead, a range of possible brightnesses is considered, and the “best” brightness is selected as the single MT brightness that minimizes a specific cost function, described below. Possible brightness values are linearly spaced between 5% of the maximum intensity value of the green signal to the maximum value of the green signal. This range assumes that there will be between one and twenty MTs in the neuron, a reasonable assumption based on previously published data (Chalfie and Thomson, 1979). Twenty is considered the upper bound to accelerate computation, but for more complex neurons, smaller brightness values can also be considered. The success of this grid search optimization is validated in the Results section.

The cost function *C* to be minimized relies on the previously asserted assumption that a change in the level of the line scan quantized by a single MT brightness marks the beginning or the end of a MT, referred to respectively as a step up or a step down. In Supp. Fig. 3(a), the black dots represent the MT starts in the quantized signal. As an initial, naive cost, we calculate the difference between the number of black dots and the number of steps up we expect to see, *N*_*exp*_. *N*_*exp*_ is the total number of MTs identified from the red line scan, *N*, less the expected number of steps up that are hidden by overlapping MT ends, or steps down. These steps down are expected to occur uniformly along the signal. If there are *N* MTs, some of the *N* steps up may coincide with steps down and not be visible in the quantized signal, i.e., there may be red triangles without corresponding black dots. For *N* MTs and a signal of length *d*, the expected fraction of pixels containing a step down is *N*/*d*, because there are *N* steps down uniformly distributed along *d*. The expected number of steps up that overlap with a step down is (*N*/*D*) × *N*, since the steps up are also assumed to be uniformly distributed along the signal.

To improve the performance of this initial cost, we can calculate additional costs which incorporate the additional information collected, namely the locations of the MT minus-ends and the green microtubule intensity. Ultimately, *C* comprises the weighted sum of three costs. Supp. Fig. 3 shows example signals and the three costs for one animal, as well as the total cost.

The second cost is the number of steps up in the quantized signal which do not colocalize with a red minus-end. To allow a buffer for effects of blurring or noise, this cost is the number of black dots in Supp. Fig. 3(b) which are more than three pixels away from the red triangles. The final cost is the mean error between the quantized signal and the observed intensity, i.e., the error between green line scan and black quantized signal in Supp. Fig. 3(c). The best single MT brightness is the brightness that minimizes this energy function. See Algorithm 3 for mathematical descriptions of these costs. Together, these three costs encourage a single MT brightness that matches the original signal well but also avoids introducing significantly more MTs than there are minus-ends. The weights of the three costs were selected to best match the various ground truth data sets available, which are described in more detail in the Results section.

### 3.3 Organization Parameter Calculation

Once we have obtained the single MT brightness, we calculate the organization parameters. This calculation is summarized in Supp. Table 1. The table defines each parameter using individual microtubules, real example signals of the entire MT bundle, and specific equations.

Of the three organization parameters calculated by MTQuant, the most straight-forward is spacing. Spacing for an individual worm is simply defined as the average number of microns between each microtubule starting location, as shown in the first row of Supp. Table 1. Let *S*_*i*_ be the distance between the *i*^*th*^ minus-end and the (*i* − 1)^*th*^ end. Alternatively, let the set *R* represent the locations of each peak identified in the red signal. *N* denotes the number of microtubules identified and *d* the length of the signal. Then, the spacing can be defined as:

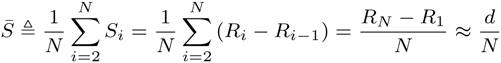

To calculate the coverage, first consider the green intensity signal, *I*_*G*_(*x*), quantized by the single MT brightness, *b*\*, to yield the quantized signal *Q*_*b*\*_(*x*), shown in the second row of Supp. Table 1. Given this quantized signal, the coverage at any location along the neuron is simply the value of *Q*_*b*\*_(*x*). Since we are calculating the average coverage, we can eliminate quantization errors by integrating the observed green intensity *I*_*G*_(*x*) instead of the quantized signal. This integration is more accurate because the area under the curves is preserved by the blurring, and the noise is zero-mean, so integrating the intensity directly causes the noise to average to zero. Therefore:

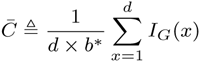

#### Algorithm 3: Extract Single MT Brightness

1. Trace and preprocess the green intensity, *I*_*G*_(*x*), and the red intensity, *I*_*R*_(*x*)(Section 3.1)
2. Identify *R* from *I*_*R*_(*x*), the set of MT minus-end locations (Section 3.1.3) *N* = ∥*R*∥_0_, the size of the set *R*
3. Calculate all possible single MT brightnesses, *B*, which are 1000 linearly spaced samples between *max*(*I*_*G*_(*x*))/20 and *max*(*I*_*G*_(*x*))
4. For each possible brightness *b* in *B*,
  a. Quantize *I*_*G*_(*x*) to generate *Q*_*b*_(*x*) = *round*(*I*_*G*_(*x*)/*b*)
  b. Identify locations of steps up in *Q*_*b*_(*x*), *U*
  c. Calculate the three costs
    1. Difference in number of steps up in *Q*_*b*_(*x*) and expected number of steps up based on red signal:

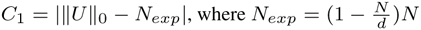
    2. Number of steps up in *Q*_*b*_(*x*) that do not colocalize with any MT minus-ends:

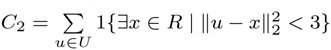
    3. Mean squared error between observed signal *I*_*G*_(*x*) and quantized signal *Q*_*b*_(*x*)

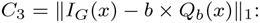
    4. Calculate total cost *C*(*b*) = *α*_1_*C*_*1*_ + *α*_2_*C*_2_ + *α*_3_*C*_3_
  d. Select 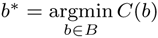

Since we cannot identify individual MTs, we cannot explicitly calculate the length of each microtubule. However, we can calculate the average length, again by integrating the green intensity. If we again begin with the quantized signal, the value of *Q*_*b*\*_(*x*) at every discrete pixel *x* is effectively *Q*_*b*\*_(*x*) pixel-wide microtubule segments at location *x*. Then if we integrate the area under the curve, as in the third row of Supp.Table 1, we have accumulated all of the one-pixel-wide microtubule segments, of all of the “microtubule material,” in the neuron. Then, we can divide by the total number of microtubules to calculate the average length of “microtubule material” in each microtubule. Again, integrating over the original intensity signal is more accurate due to the blurring and additive noise corrupting the signal.

Since the MT minus-ends are distributed along the entire length of the axon, we can approximate that the last minus-end is close to the end of the signal, i.e., *R*_*N*_ ≈ *d*. Therefore the length is approximately equal to the product of the coverage and the spacing.

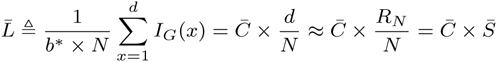

This relationship between 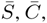, and 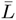 highlights that there are in fact only two independent organization parameters. We consider all three in our analysis, however, due to the varying intuitive insights each of the three parameters provides.

### 3.4 Distribution Refinement

The organization parameters calculated thus far are in reality noisy measurements of the true, hidden organization parameters. This noise is a result of imaging noise, biological noise, and calculation error. The organization parameters are expected to have some variation across a particular group of animals, as defined by age or genotype, and in order to differentiate subtly different groups, this additional noise must be somehow mitigated. The observed organization parameters are directly related to the hidden parameters. We assume a mixture of Gaussians model; given the value of the hidden parameters, the observed parameters are distributed normally. This mixture of Gaussians model can be solved using expectation-maximization (EM), which aggregates the observed organization parameters from all of the animals to refine the estimates of the distributions of average spacing, length, and coverage over a population (Friedman *et al.,* 2001).

The EM algorithm iteratively identifies the probability distributions of unobserved variables from observed data. Here, we observe the organization parameters as the result of the individual analysis: the average spacing, 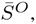, length, 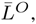, and coverage, 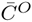 of the MTs for that worm. The EM algorithm alternately calculates conditional probability distributions of 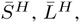 and 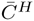 given the observed data, and then calculates the mean and variance of the probability distributions of the parameters that best generate these conditional distributions.

The application of EM is the same for each of the three parameters. As an example, consider the refinement of the distribution of 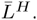. For each animal *i*, and each possible average length 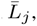, the conditional probability distribution 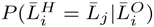 is updated using Bayes rule:

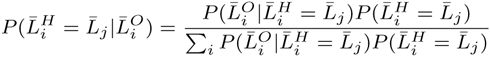

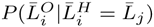 is assumed to be Gaussian, based on observed data. 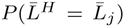 is initialized with the frequency histogram of the initial observed values 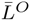, and then updated during the refinement process. It is straightforward to calculate 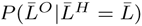 and to update the marginal distribution 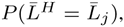, because this is a mixture of Gaussians model (Friedman *et al*., 2001). These steps are repeated for 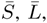 and 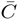 until 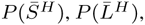, and 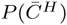 converge. In general, EM has been shown to converge (Wu, 1983). After applying EM, the average value of each parameter across a population of worms does not change, but the variance decreases, as shown in Supp. Fig. 4.

### 3.5 Modeling

In order to verify the accuracy of the MTQuant framework, it was applied to artificial green tubulin and red patronin signals that simulated the real data. The point spread function and the periodogram of the noise of the imaging system were measured with 100nm tetra-speck fluorescent beads (Life Technologies T-7279) and are plotted in Supp. Fig. 5(a)-(b). We approximate the PSF as a Gaussian Zhang *et al.* (2007). To simulate the signals, inter-arrival distances of the starting locations of MTs were drawn from an exponential distribution to correspond to a Poisson process. The mean of the distribution was adjusted to match the observed number of red dots identified in the red fluorescence images. For each MT, a length was randomly generated using a previously identified empirical distribution (Yu and Baas, 1994). These MTs were summed to form the quantized green signal. The quantized signal was scaled by a randomly generated step size to simulate the intensities of the fluorophores, which vary from animal to animal due to biological variation of gene expression and imaging conditions. To generate the red signal, at each generated MT starting location, a box of random width and height was placed to represent the variable amount of patronin protein that can collect at the tip of a MT (Supp. Fig. 5(c)-(d)). Then both signals were blurred according to the measured point spread function of the imaging system and noise was added to the signals that mimicked the noise measured in the imaging system (see Supp. Fig. 5(e)-(f)). To generate the noise, a white Gaussian noise signal was passed through a linear filter with the frequency response of the square root of the measured noise periodogram.

## 4 Results

MTQuant has been verified both numerically and biologically. The various verification methods, described below, showed satisfactory consistency between the ground truth and the calculated results. Armed with this verified method, we first studied the relationship of animal age to microtubule organization, presented here.

### 4.1 Mathematical Verification

MTQuant was first applied to simulated data in order to verify the algorithm mathematically. Simulated, quantized signals were corrupted with blurring and noise, as described in Supp. Fig. 5(a)-(f) and Section 3.5. Then, the simulated intensity signals were processed with the framework, and the error in single MT brightness was reported. Fig. 2(a) shows the calculated brightness for 100 trials plotted against the model brightness for the same simulated signals. Without knowledge of the blurring or the noise of the imaging system, the algorithm successfully calculated the single MT brightness for the simulated data (p-value = 1.56e-50).

**Fig. 2.**
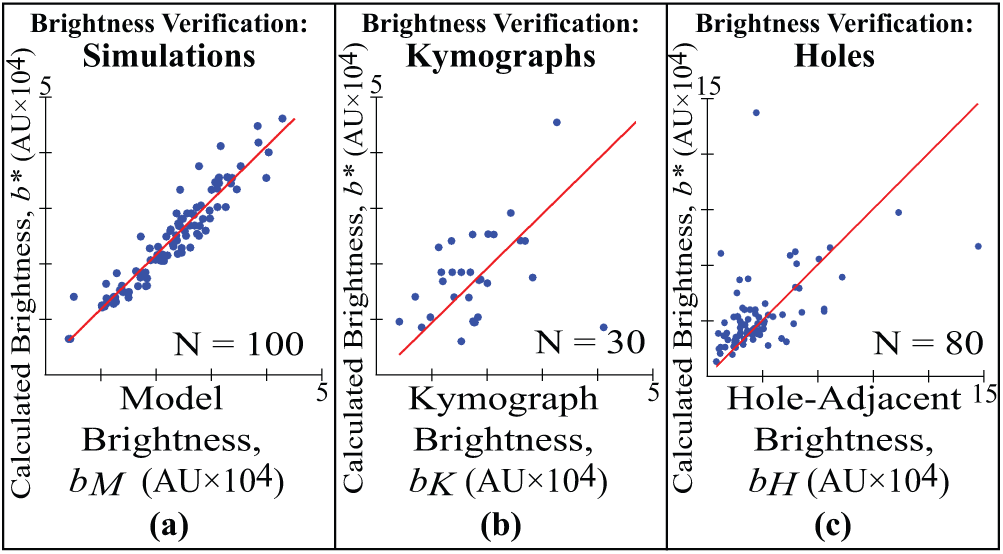
MTQuant verification results. (a) Single MT brightnesses calculated with MTQuant, *b*\*, and from model, *b*_*M*_ (p-value = 1.56e-50). Red line is *b*\* = *b*_*M*_. (b) *b*\* versus single MT brightnesses from kymographs, *b*_*k*_ (p-value = 0.038). Red line is *b*\* = *b*_*k*_. (c) *b*\* versus single MT brightnesses around holes, *b*_*H*_ (p-value = 5.78e-07). Red line is *b*\* = *b*_*H*_.

### 4.2 Biological Verification

The most basic biological verification compared the spacing, coverage, and length of MTs obtained via serial reconstructions of electron microscopy sections to the distributions calculated using MTQuant. Both the previously published coverage and length of MTs in (Chalfie and Thomson, 1979), and the more recently calculated organization parameters obtained in (Yogev *et al*., 2016b), align well with our distributions of MTs in wild-type worms.

To show that MTQuant can enable biologists to compare groups of animals, the framework was next applied to two sets of fluorescence images. MTQuant was first applied to kymographs of MTs. Kymographs are images of the tubulin intensity change over time. In Supp. Fig. 5(g)(i), the x-axis denotes position, and the y-axis denotes time. The solid lines in Supp. Fig. 5(g)(ii) trace a single MT growing and shrinking. The brightness of a single MT can be calculated as the difference in the average intensities of the dotted lines above and below each solid line. Since DA9 has relatively few neurons, and they are fairly stable, it is reasonable to assume that almost all intensity changes reflect the dynamic behavior of a single MT. To validate this assumption, we measured MT dynamics with an independent method, using the end-binding protein EBP-2::GFP, and found that it was low (Yogev *et al*., 2016b).

Before calculating the average single MT brightness for each kymograph, we must correct for photobleaching. Fluorophore intensities diminish over time due to a phenomenon called photobleaching (Vicente *et al*., 2007). The average intensity of each row is assumed to drop as a bi-exponential, i.e., *y*(*x*)= *αe*^*βx*^ + *γe*^*δx*^ where *x* is the row number, and *y*(*x*) is the average intensity of all pixels in that row, scaled to have a maximum value of 1. We use MATLAB’s `fit` function to solve for *α*, *β*, *γ*, and *δ*. Then we scale the intensities in each row *x* by 1/*y*(*x*). As a result, pixels toward the bottom of the image become brighter, but intensities at the top of the image do not change significantly.

Fig. 2(b) shows the single MT brightness calculated using MTQuant and the average brightness change around 369 growing or shrinking MTs in 30 kymographs. There was a clear correlation between the two sets of measurements (p-value = 0.038), so relative changes in brightness across animals can be detected, and, in turn, so can relative changes in coverage and length.

A set of “axon hole” images was used as the second collection of ground truth images. In these images, there was a gap in the continuity of the MT coverage of the axon, as in Fig. 2(c). Such gaps are relatively rare, and they occur mostly in younger animals and in portions of the axon that are far from the cell body. The average intensity change on either side of the holes typically corresponds to the brightness of a single MT, although we cannot exclude the possibility that two MTs end simultaneously. Fig. 2(c) compares the single MT brightness calculated by MTQuant and the intensity measured in the images around 184 holes that were selected manually in 80 images. There is a strong correlation between the two sets of brightnesses (p-value = 5.78e-07).

### 4.3 Biological Results

As an initial experiment, the organization parameters of wild-type worm MTs were calculated at different ages. Fig. 3 shows the organization parameters as worms age from 12 hours to one week. Fig. 3(a) shows that, as worms age, the average spacing between MTs does not change significantly (p-value = 0.053). However, Fig. 3(b)-(c) do show significant changes. The length of MTs increases as worms develop (p-value = 1.46e-74), consistent with EM studies (Yu and Baas, 1994; Chalfie and Thomson, 1982). Consequently, the average coverage of MTs also increases as a function of worm age (p-value = 4.89e-60). The significance of these relationships was calculated using Spearman rank correlation (McDonald, 2014).

**Fig. 3.**
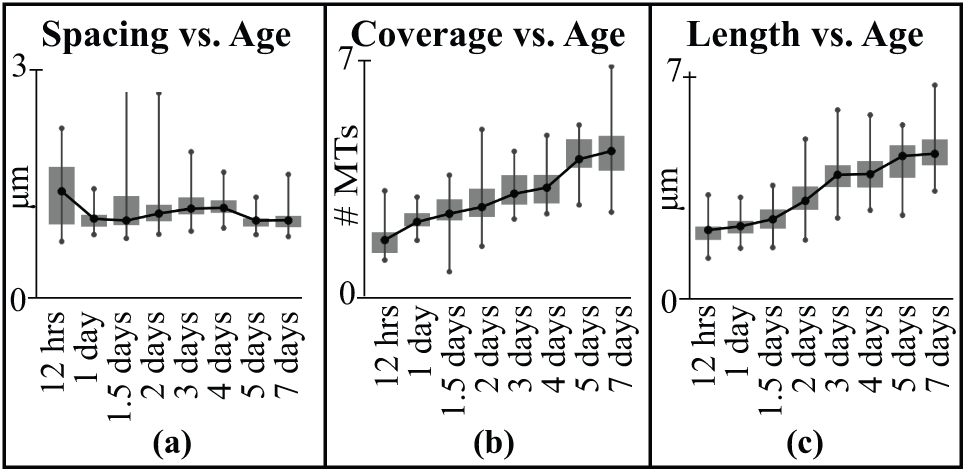
Microtubule Organization as a function of animal age. (a) Spacing of MTs does not change significantly from worm larvae to adult worms (p-value = 0.053). (b) Coverage, or number of MTs in axon cross-section, significantly increases with age (p-value = 4.89e-60). (c) Average MT length also increases significantly with age (p-value = 1.46e-74). All p-values calculated using Spearman rank correlation.

The increases in MT length and coverage suggest that, as the axon grows, existing MTs also elongate, and new MTs develop to tile the axon. This important result has implications as to how neurons elongate and how the cytoskeleton scales during development.

This result demonstrates the utility of MTQuant. The relationship between age and MT organization would have been nigh impossible to achieve without this rapid, automated method. We have also applied MTQuant to over 3000 animals to ask a variety of other biological questions about microtubules, delineated in (Yogev *et al*., 2016b).

## 5 Conclusion

The MTQuant framework provides a rapid and robust method of quantifying MT organization, allowing biologists to study the spatial distribution of MTs on a large scale in vivo using light microscopy instead of more labor-intensive and error-prone methods such as serial reconstructions of electron microscopy. Even super-resolution imaging, which requires highly specialized equipment and biological markers, does not provide sufficient resolution to visualize individual MTs. Instead, MTQuant allows biologists to “see” information about MT organization that is significantly below the resolution limit of light microscopy.

MTQuant has been validated on three ground truth data sets. The organization parameters of wild-type worms calculated by MTQuant also match previously published data (Chalfie and Thomson, 1979). MTQuant was applied to hundreds of MT images of animals at varying ages. MT spacing in wild-type animals remained constant as the axon grew over three-fold during during development. However, MTs elongated during development, suggesting that as animals grow, existing MTs elongate, and new MTs populate the new portions of the axon.

MTQuant has been used to answer various biological questions (Yogev *et al*., 2016b), and can be used more specific, subtle MT organization studies as well: various sections of the dendrite have been studied separately using MTQuant, to quantify the MT material that appears to cluster in the tip of the dendrite in some mutants (Yogev *et al*., 2016a).

By exploiting the fact that MTs form a bundle, we were able to extract a significant amount of information from the fluorescent images of MTs. Since this framework makes few assumptions about MTs beyond their bundled organization, MTQuant can potentially be applied to the study of other filaments as well, such as intermediate filaments, which are organized similarly to MTs (Chang and Goldman, 2004). Other potential applications include the quantification of myelin wraps which also form quantized steps along the axon (Snaidero *et al*., 2014). In general, MTQuant can be extended to discrete structures so long as their tagged intensities are expected to form quantized steps.

For biological data with low signal-to-noise ratios, combining multiple measurements is essential for achieving high quality outcomes. Aggregating results from multiple animals when extracting the organization parameters yields more accurate estimates of the distributions of the underlying biological data. By incorporating simple assumptions about MT organization with multiple measurements, this framework has enabled us to gain a deeper understanding of MTs in *C. elegans.*

## Acknowledgments

We thank Jing Xiong for her suggestion to use a mixture of Gaussians model for the Distribution Refinement block.

## Supplementary Figures

**Supplementary Figure 1:**
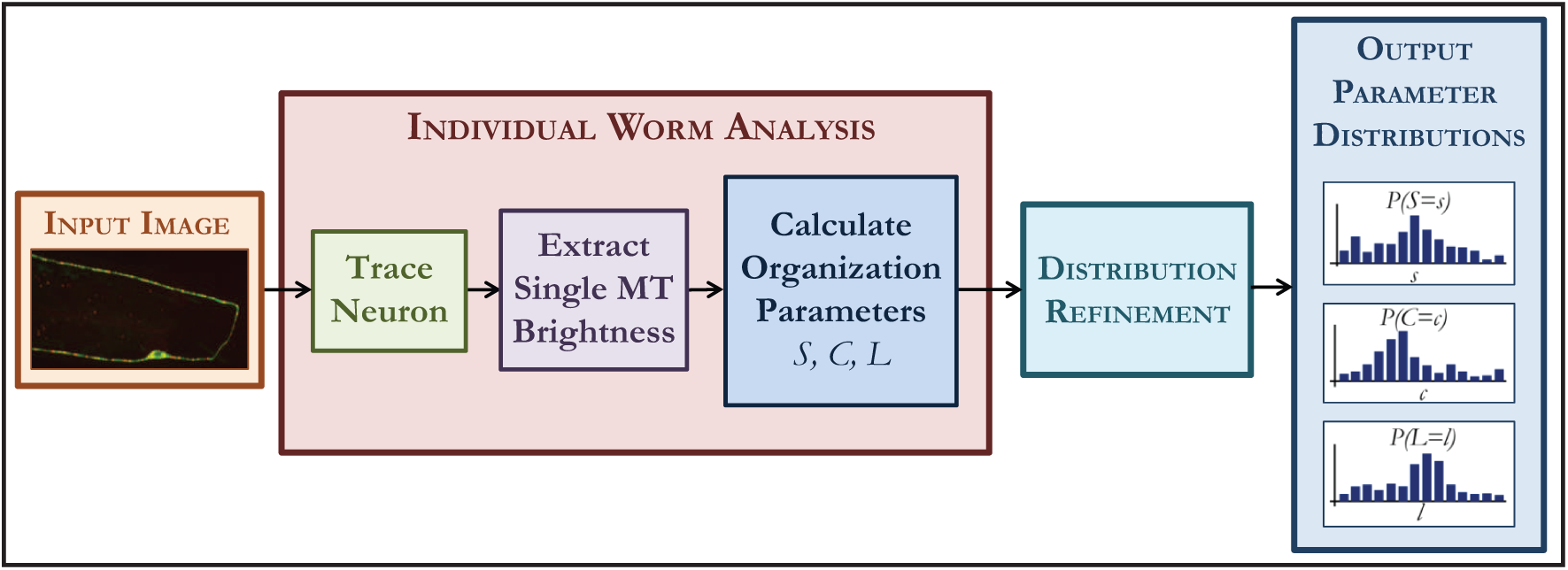
MTQuant Block Diagram: individual worm analysis followed by distribution refinement.

**Supplementary Table 1:**
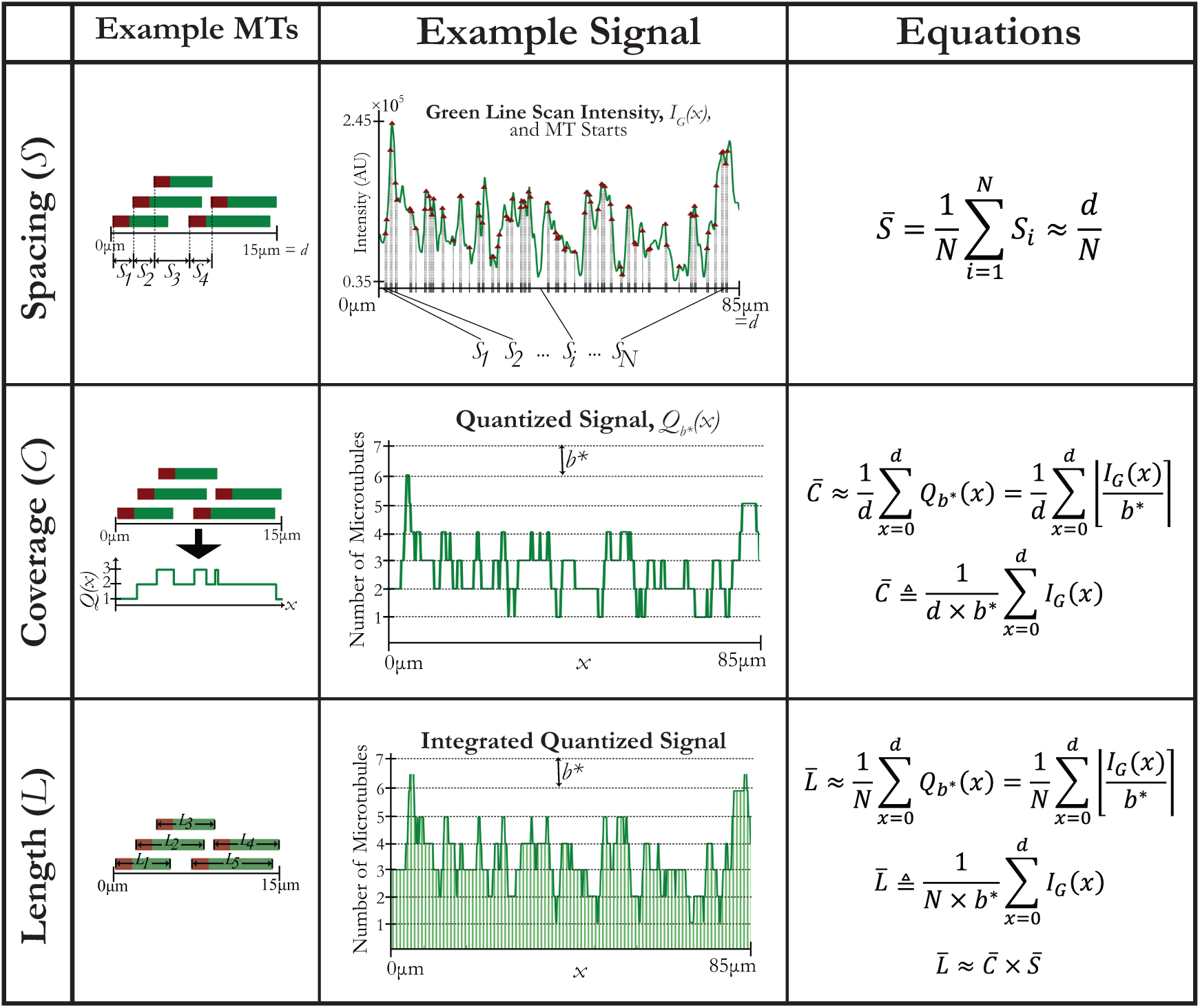
Organization Parameter Calculation. Each panel contains a theoretical schematic (i), an example signal (ii), and equations for calculation (iii). (a) Spacing, 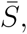 is average microns between MT starts, or approximately signal length, *d,* divided by total number of MT starts, *N*. (b) Coverage, 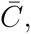 refers to average number of MTs in bundle cross-section, or average value of quantized signal, *Q*_*b*\*_(*x*), along neuron. To compensate for blurring and noise, a more accurate 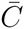 is the integral of green signal *I*_*G*_(*x*) divided by *d* and single MT brightness, *b*\*. (c) Length of each MT cannot be calculated since we cannot identify individual MTs as in (i), but we calculate average MT length, 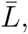 by integrating *I*_*G*_(*x*) and dividing by *N* and *b*\*. Organization parameters are not all independent; 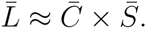

|  | Example MTs | Example Signal | Equations |
| --- | --- | --- | --- |
| Spacing ($\bar{S}$) | | | $\bar{S} = \frac{1}{N} \sum_{i=1}^N S_i \approx \frac{d}{N}$ |
| Coverage ($\bar{C}$) | | | $\bar{C} \approx \frac{1}{d} \sum_{x=0}^d Q_{b^*}(x) = \frac{1}{d} \sum_{x=0}^d \left[ \frac{I_G(x)}{b^*} \right]$ $\bar{C} \triangleq \frac{1}{d \times b^*} \sum_{x=0}^d I_G(x)$ |
| Length ($\bar{L}$) | | | $\bar{L} \approx \frac{1}{N} \sum_{x=0}^d Q_{b^*}(x) = \frac{1}{N} \sum_{x=0}^d \left[ \frac{I_G(x)}{b^*} \right]$ $\bar{L} \triangleq \frac{1}{N \times b^*} \sum_{x=0}^d I_G(x)$ $\bar{L} \approx \bar{C} \times \bar{S}$ |

**Supplementary Figure 2:**
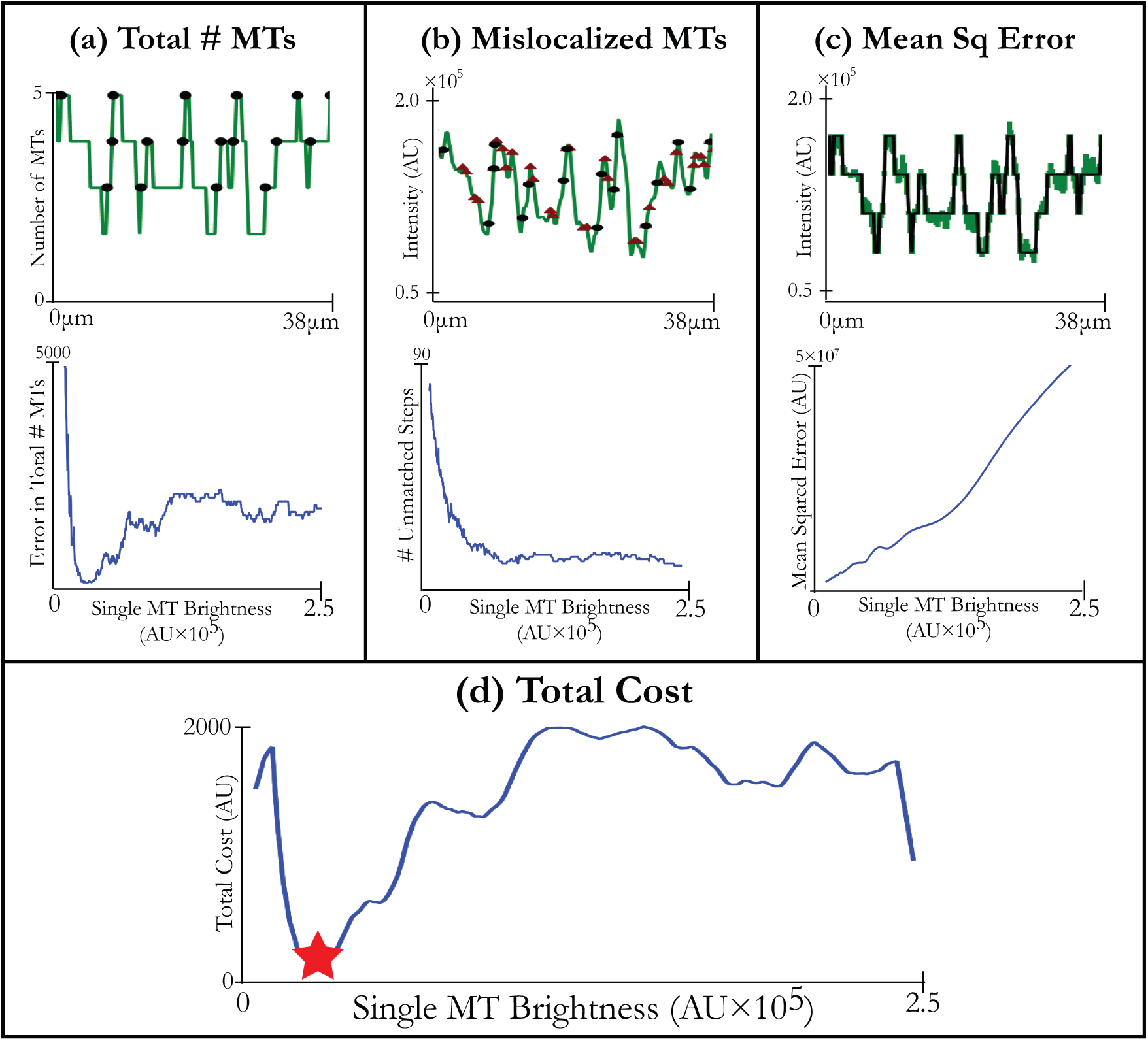
Cost calculation for single MT brightness. For each possible single MT brightness, green signal is quantized and compared to the original signal. In (i)-(iii), top row represents an example signal, and bottom row displays corresponding cost. (i) Squared error between number of steps up in the quantized signal (black dots) versus the expected number of steps based on the observed number of patronin puncta. (ii) The number of steps up in the quantized signal (black dots) that are more than 3 pixels away from a patronin puncta (red triangles). (iii) Mean squared error between quantized signal and green tubulin signal. (iv) Total cost. Red star corresponds to best single MT brightness, *b*\*.

**Supplementary Figure 3:**
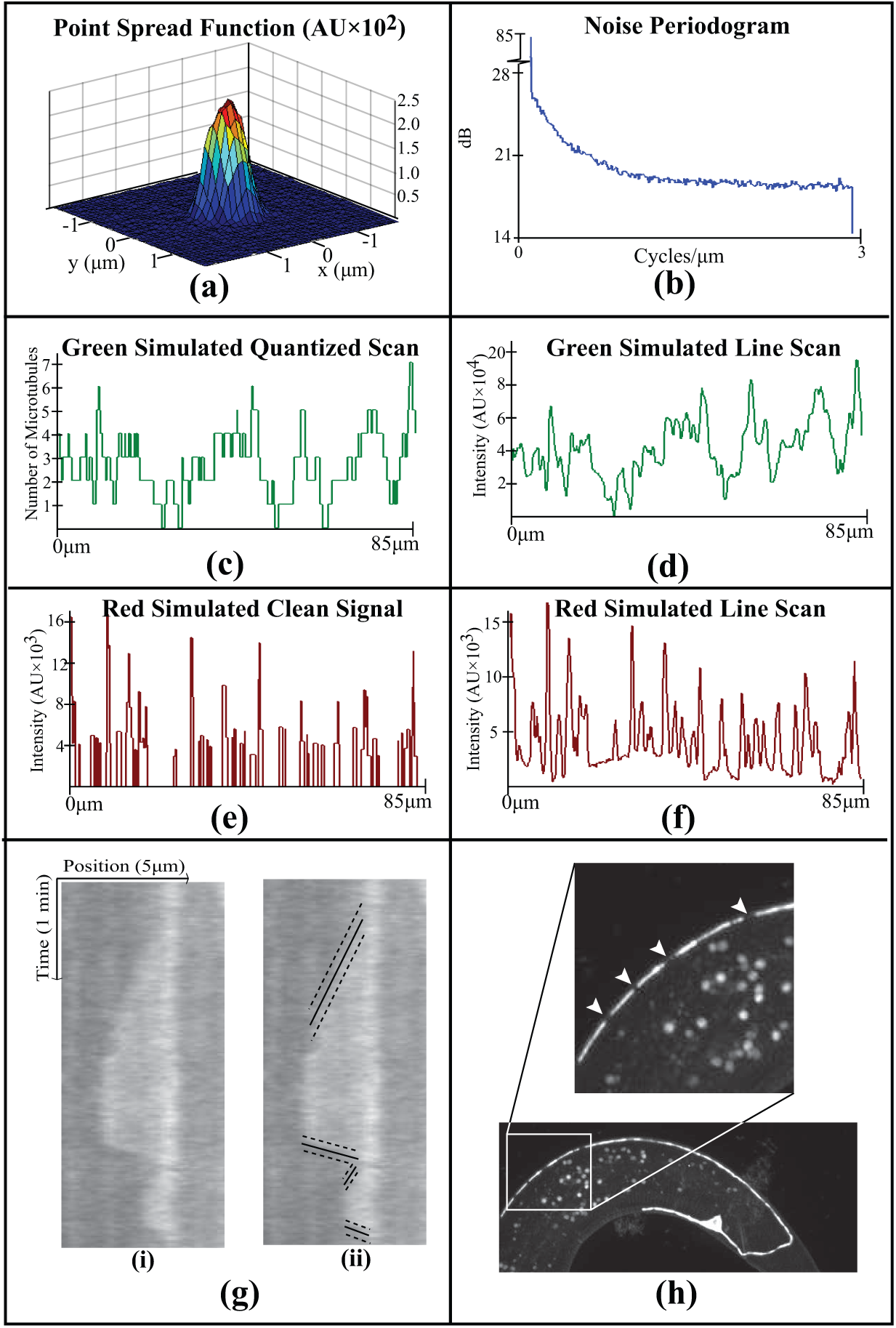
MT verification. (a) Measured point spread function of imaging system. (b) Measured periodogram of noise profile of imaging system. (c) Simulated clean tubulin signal. (d) Tubulin signal with blurring and noise. (e) Clean simulated patronin signal. (f) Patronin signal with burring and noise. (g) Example kymograph of MT growing and shrinking (solid line). The single MT brightness is the different in the average intensities along pairs of dotted lines. (h) Example of holes in MT bundle (arrowheads in inset).

